# Lgr5+ telocytes are a signaling hub at the intestinal villus tip

**DOI:** 10.1101/850909

**Authors:** Keren Bahar Halpern, Hassan Massalha, Rachel K. Zwick, Andreas E. Moor, David Castillo-Azofeifa, Milena Rozenberg, Lydia Farack, Adi Egozi, Dan R. Miller, Inna Averbukh, Yotam Harnik, Noa Weinberg-Corem, Frederic J. de Sauvage, Ido Amit, Ophir D. Klein, Michal Shoshkes-Carmel, Shalev Itzkovitz

## Abstract

The intestinal epithelium is a structured organ composed of crypts harboring Lgr5+ stem cells, and villi harboring differentiated cells. Spatial transcriptomics have demonstrated profound zonation of epithelial gene expression along the villus axis, but the mechanisms shaping this spatial variability are unknown. Here, we combined laser capture micro-dissection and single cell RNA sequencing to uncover spatially zonated populations of mesenchymal cells along the crypt-villus axis. These included villus tip telocytes (VTTs) that express Lgr5, a gene previously considered a specific crypt epithelial stem cell marker. VTTs are elongated cells that line the villus tip epithelium and signal through Bmp morphogens and the non-canonical Wnt5a ligand. Their ablation strongly perturbs the zonation of enterocyte genes induced at the villus tip. Our study provides a spatially-resolved cell atlas of the small intestinal stroma and exposes Lgr5+ villus tip telocytes as regulators of the epithelial spatial expression programs along the villus axis.

## Introduction

The small intestine is a structured organ composed of repeating crypt-villus units. Lgr5+ epithelial stem cells at the base of the crypts continuously divide to give rise to proliferative progenitors that migrate towards the villi^1,2^. As they approach the crypt exits, these progenitors become post-mitotic and differentiate into distinct lineages – absorptive enterocytes, mucus-secreting goblet cells, enteroendocrine cells and tuft cells. The differentiated cells operate for about three days as they continue to migrate along the villus walls until they are shed from the villi tips^3^.

Villi cells have traditionally been considered ‘terminally-differentiated’ in the sense that they have irreversibly committed to a functional cell state, which they carry from birth to death. Recent studies challenged this view and demonstrated that villi epithelial cells constantly change their functional state, such that around 85% of enterocyte genes are expressed in a non-uniform manner along the villus axis^4^. Enterocytes first implement anti-microbial programs at the base of the villi, then shift to sequential absorption of amino acids, carbohydrates and lipids in distinct villi zones, finally up-regulating genes associated with cell adhesion and immune modulation at the villi tips. Similarly, enteroendocrine cells change the types of hormones they produce as a function of their position along the villus axis^5,6^. The epithelial lineages along the intestinal villus therefore exhibit profound spatial heterogeneity.

What are the mechanisms that facilitate the spatial heterogeneity of the villus epithelium? One mechanism could be an internal ‘clock’, whereby epithelial cells are pre-programmed to turn distinct functions on and off at different times. A second mechanism could be a zonated response to spatial gradients of nutrients and bacteria at the luminal sides of the tissue. Yet a third mechanism could entail zonated molecular cues that originate in the lamina propria, the underlying stroma at the basal sides of the epithelial layer^7–11^. Here, we use spatially-resolved transcriptomics to characterize the zonated stromal gene expression signatures along the crypt-villus axis. We identify four mesenchymal cell populations residing at distinct crypt-villus zones. These include a sub-population of telocytes localized at the villus tip that is marked by Lgr5, a gene previously considere a specific marker of epithelial crypt stem cells. We demonstrate that Lgr5+ villus tip telocytes regulate the epithelial gene expression programs at the villus tip.

## Results

### A spatial expression atlas of the small intestinal stroma

To obtain a global view of spatial heterogeneity of stromal gene expression we used Laser Capture Microdissection (LCM) to isolate four stromal zones along the jejunum crypt-villus axis from five mice (Fig. 1a,b). We performed RNA sequencing of these segments, yielding a coarse zonation map (Table S1). We focused on the zonation patterns of ligands and receptors^12,13^, since these are likely to implement zonated cross-talk with the epithelium (Fig. 1c, Extended Data Fig. 1a,b, Table S2). Examples of ligand/receptor genes that exhibited higher expression in the crypt stroma included Grem1, encoding the BMP pathway inhibitor Gremlin1^14^ (Fig. 1c) and Il18r1, the receptor for Il18. The cytokine Il18 has been shown to be expressed in enterocytes at the lower villi zones, as part of an anti-microbial gene module^4^. Using single molecule Fluorescense in-situ Hybridizatoin (smFISH^15^), we identified type 3 innate lymphoid cells (ILC3) expressing Il18r1 at the stromal side of these zones (Extended Data Fig. 1c-g), suggesting spatial recruitment of these immune cell components by zonated enterocyte Il18 signal.

**Fig. 1.**
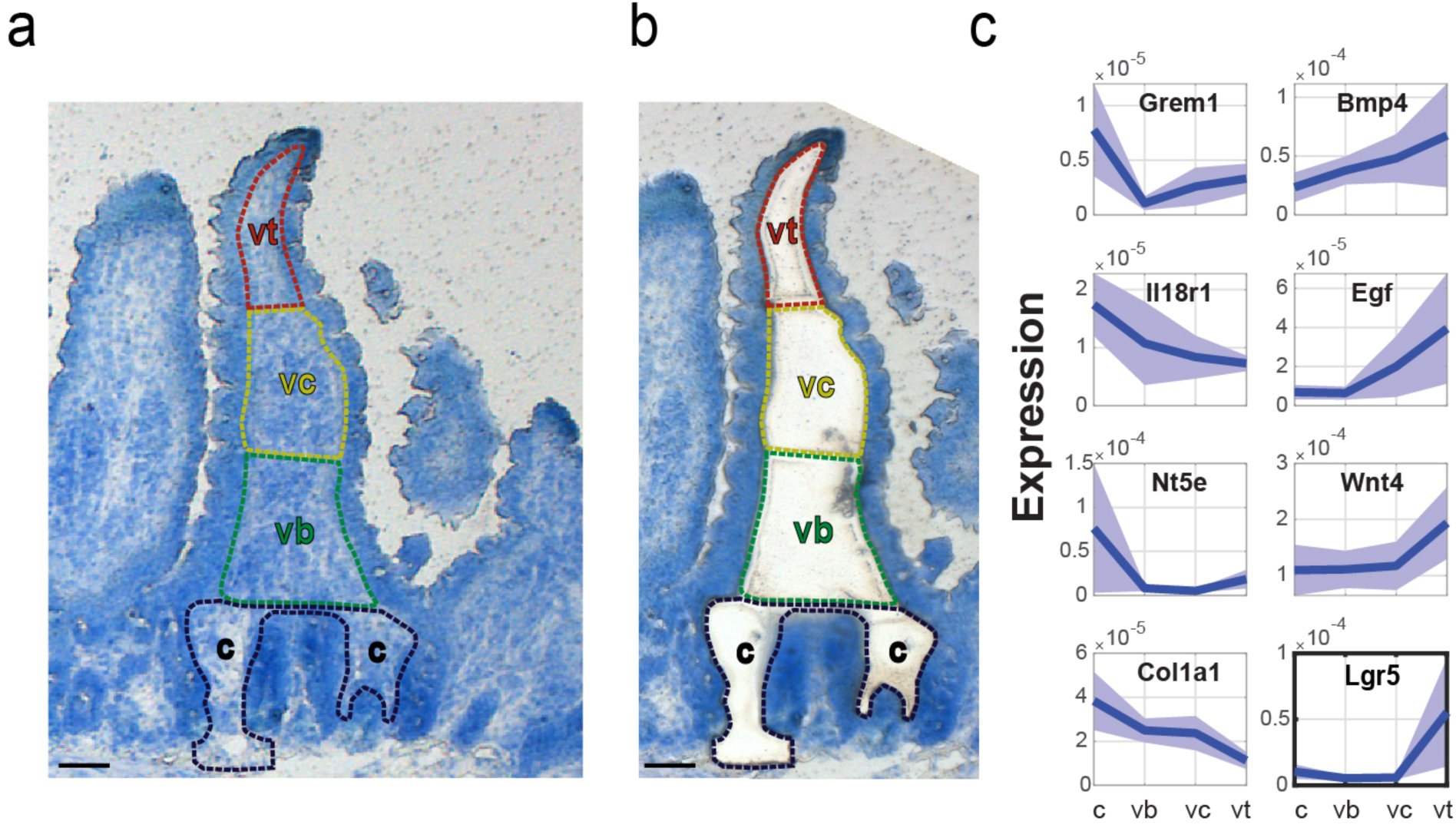
Spatial transcriptomics of the intestinal stroma. a-b) LCM of four zones along the crypt-villus axis before (a) and after (b) laser dissection. c – crypt, vb – villus bottom, vc – villus center, vt – villus tip. Scale bar – 50 µm. c) Representative spatial LCMseq expression profiles of stromal ligands and receptors zonated towards the crypt (left) or villus tip (right). Units are fraction of sample mRNA, patches are SEM.

### Lgr5 is abundantly expressed in stromal cells at the villus tip

We next focused on stromal ligands and receptors that were zonated towards the villi tips in our stromal LCM-RNAseq data. These included Bmp4, a morphogen that was shown to inhibit epithelial proliferation^16^ and to control enteroendocrine cell differentiation pathways at the tip^5^ (Fig. 1c). Other villus tip molecules included the non-canonical Wnt ligand Wnt4 and the epidermal growth factor Egf. Egfr, the receptor for Egf, has elevated expression levels in villus tip enterocytes^4^, constituting another example of correlated spatial expression of pairs of epithelial and stromal ligands and their matching receptors (Table S2). We further analyzed zonated proteins constituting the extracellular matrix (ECM, also termed the ‘matrisome’^17^), identifying distinct collagens and matrix metalloproteinases that are differentially zonated between the crypt and villus stroma (Extended Data Fig. 1h). These included the crypt enriched Lama2, encoding the laminin subunit alpha-2 and the villus enriched Lama5, encoding the laminin subunit alpha-5, previously shown to be differentially zonated at the protein level^18,19^.

Our zonation analysis revealed Lgr5 to be one of the most highly expressed receptors in the villus tip stroma (Fig. 1c, Table S1). Lgr5 has been shown to be a specific marker of epithelial stem cells at the crypt base^2,15^. It was therefore unexpected to observe elevated expression levels of this gene at the lamina propria of the villus tip. To validate this finding, we performed smFISH and detected abundant localized expression of Lgr5 transcripts in PDGFRa+ telocytes that co-expressed Bmp4, a classic villus tip ligand^20^ (Fig. 2). Telocytes are large mesenchymal cells with elongated extensions that stain positively for the PDGFRa surface marker. They form intricate contacts with both epithelial cells and other stromal cell types^9,21^. Telocytes that surround the crypts constitute important niche cells that secrete canonical Wnt morphogens and Rspo3 to maintain stemness of crypt Lgr5+ epithelial stem cells^9,11,22,23^. The molecular identities of telocytes residing in the villi stroma have thus far not been characterized.

**Fig. 2.**
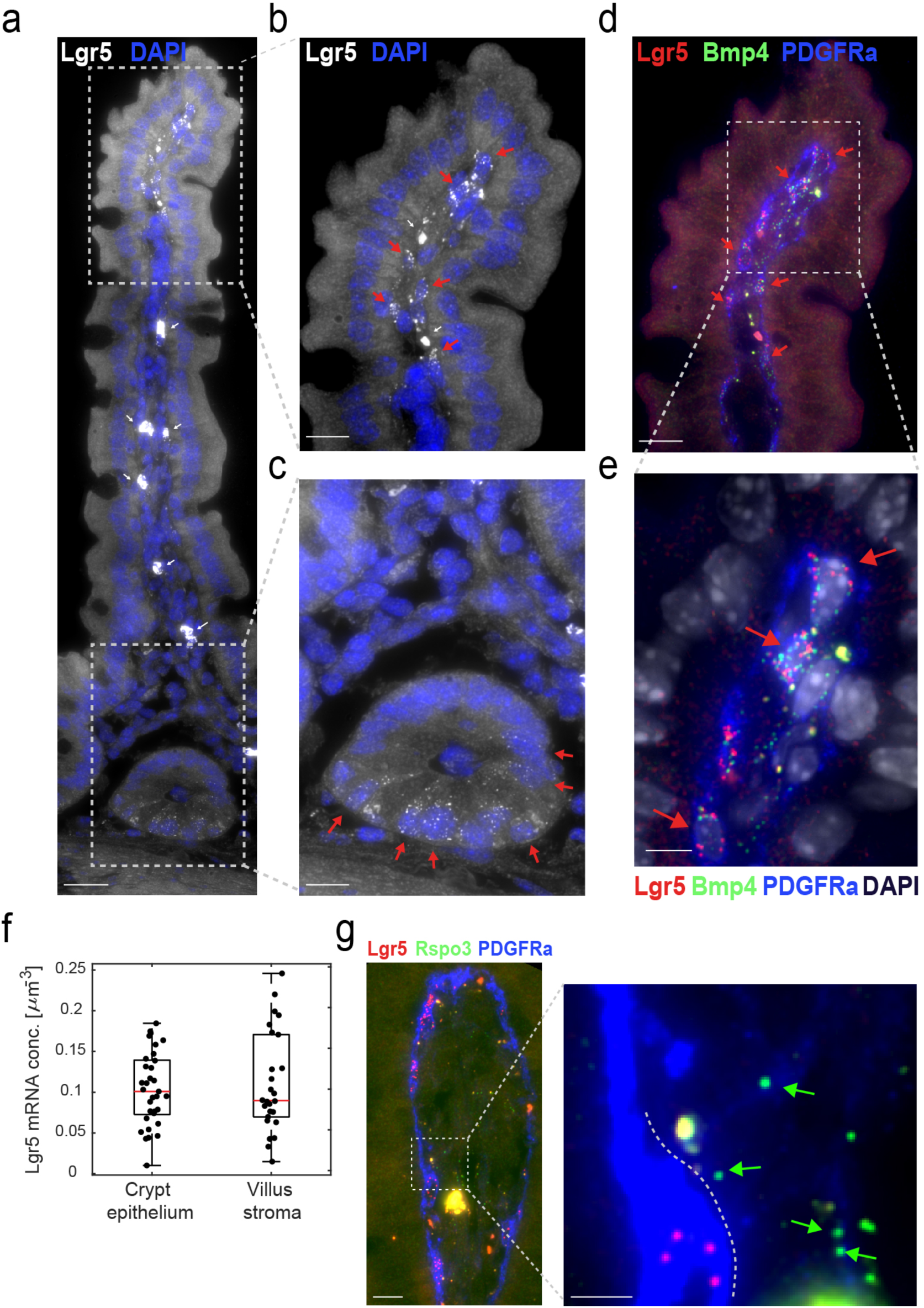
Lgr5 is expressed abundantly in villus tip telocytes. a) smFISH of Lgr5, DAPI in blue, Scale bar – 20 µm. b) Blow up of villus tip, Scale bar – 10 µm. In a,b thin white arrows point at autofluorescent blobs. c) blow up of crypt, Scale bar – 10 µm. Red arrows in b-c point to Lgr5 positive cells. d) Lgr5 mRNA (red dots) expressed in PDGFRa+ VTTs that co-express Bmp4 mRNA (green dots). Scale bar – 10 µm. Red arrows point to Lgr5 and Bmp4 double positive cells. e) Blow up of the region boxed in d). Scale bar – 5 µm. f) Lgr5 mRNA concentrations in VTTs are comparable to those in Lgr5+ crypt base columnar cells. Boxes show 25-75 percentiles of the smFISH expression, horizontal red lines are medians. g) Rspo3 mRNAs are localized on telopodes that extend away from the cell bodies of the VTTs. VTTs are marked by Lgr5 mRNA (red dots), Rspo3 mRNA (green dots) is localized away from the cell body, PDGFRa antibody mark VTTs cell bodies and telopodes. Scale bar – 10 µm, in inset, green arrows point to Rspo3 mRNAs (green dots) localized on PDGFRa telopodes (blue). Telocyte cell body is marked by white dashed line. inset Scale bar – 5 µm.

The expression levels of Lgr5 in the villus tip telocytes (VTTs) were comparable to the expression levels in the epithelial crypt base columnar stem cells (mean expression of 0.109±0.012 mRNAs/µm^-3^ in tip telocytes vs. 0.104±0.008 mRNAs/µm^-3^ in crypt stem cells, (Fig. 2f). VTTs expressed both Lgr5 and its ligand Rspo3^24^ (Fig. 2g). Notably, mRNAs of Rspo3 were localized along telopodes – thin PDGFRa+ extensions of the telocytes that extended towards the lamina propria and away from the cell body, where Lgr5 mRNAs were localized (Fig. 2g).

Several mouse models have been developed to investigate intestinal Lgr5+ crypt epithelial stem cells. These include models for ablating Lgr5 cells^25^ and for tracing their progenies^2^. We asked whether the knock-in constructs in these mice were also expressed in Lgr5+ VTTs. Indeed, VTTs were clearly seen in Lgr5–GFP-DTR mice^25^ (Fig. 3a,b) and Lgr5-EGFP-Ires-CreERT2 mice^2^ (Fig. 3c,d). All GFP+ stromal cells had Lgr5 mRNA in the Lgr5–GFP-DTR mice (60 out of 60 cells counted over 10 villi from 2 mice) and no Lgr5+ cells at the villus tip stroma lacked GFP. The proportions of Lgr5+ VTTs positive for the EGFP knock-in in the Lgr5-EGFP-Ires-CreERT2 mice were lower than the proportion of EGFP+ crypts (13%±3% vs. 35%±6%, counted over 22 villi in two mice), indicating a partial silencing of this knock-in reporter in the mesenchymal lineage in this mouse model. In summary, Lgr5 specifically labels telocytes localized at the villus tip, in addition to its documented role as a marker of epithelial crypt stem cells.

**Fig. 3.**
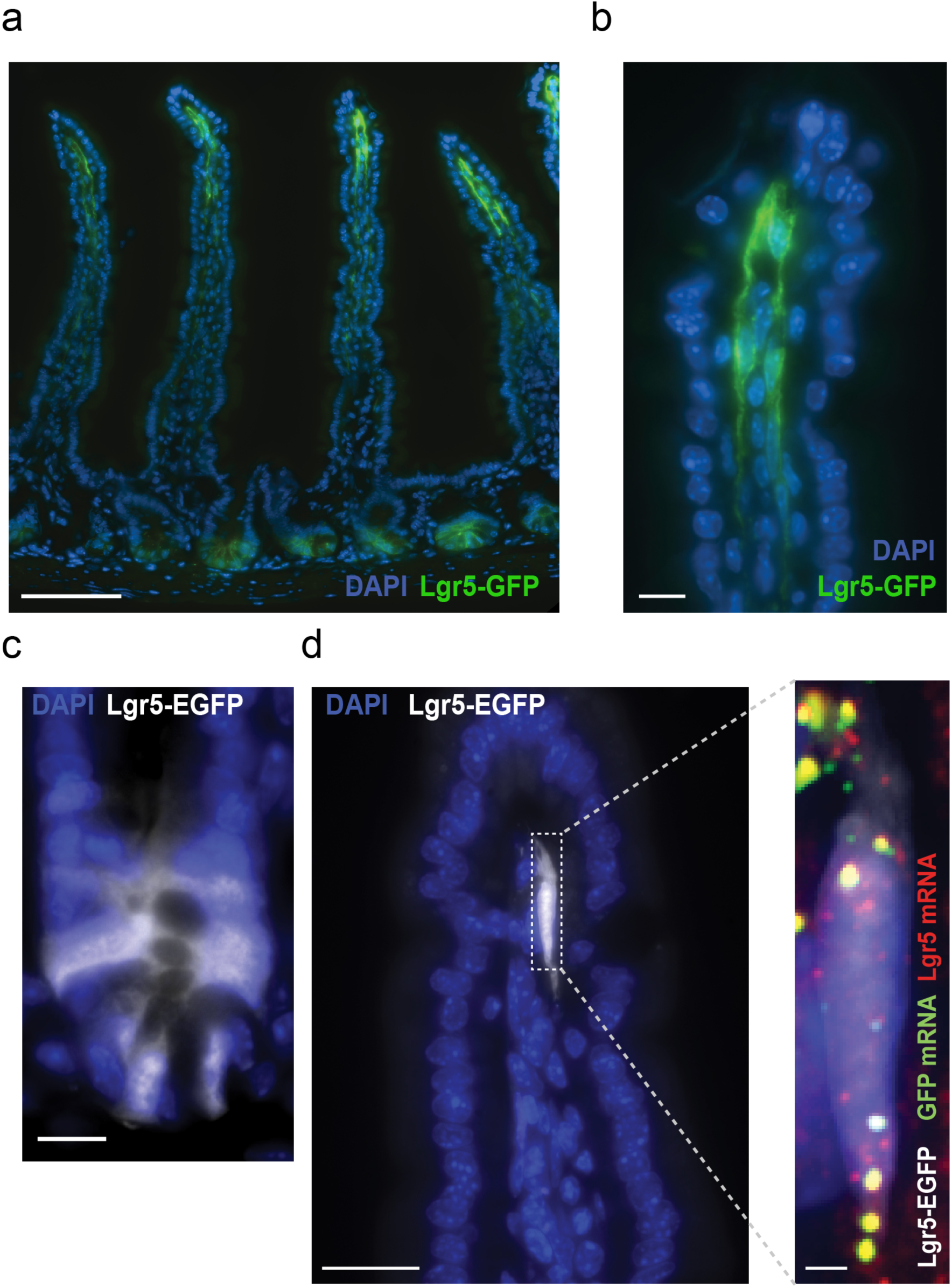
Lgr5 expression at the villus tip of Lgr5-knock-in mouse models. a) GFP fluorescence (green) observed in both crypt base columnar cells (CBCs) and VTTs in Lgr5–GFP-DTR mice. Scale bar– 100 µm. b) blow up of villus tip with GFP+ VTTs, scale bar – 10 µm. c) EGFP fluorescence of crypt base columnar cells in Lgr5-EGFP-IRES-creERT2 mice. Scale bar – 10 µm. d) EGFP fluorescence in VTTs. The Lgr5-EGFP knock-in construct is expressed in a patchy manner in the villus tip stroma. Scale bar – 20 µm. Blow up shows Lgr5 mRNAs in red, EGFP mRNA in green and DAPI in blue. Scale bar – 2 µm. Yellow blobs are autofluorescent elements.

### Single cell transcriptomics reveals the molecular signature of VTTs

We next sought to obtain the global gene expression signatures of Lgr5+ VTTs and of other intestinal mesenchymal cell populations. LCMseq provides a spatial map of gene expression along the crypt villus axis, however each zone represents a mixture of diverse cell types, including mesenchymal cells, endothelial cells, immune cells^26^ and enteric neurons^27^. Unraveling the cellular origins of the zonated transcripts requires single cell transcriptomics. Extracting telocytes is challenging due to their long thin extensions and their entrenchment within the ECM. PDGFRa was previously shown to be expressed in telocytes throughout the crypt-villus axis^9^. We therefore used fluorescence activated cell sorting to enrich for PDGFRa+ cells that included the intestinal telocytes, as well as other mesenchymal cell types (Extended Data Fig. 2). We performed single cell RNA sequencing on these cells using the MARS-seq protocol^28^. Although we enriched for PDGFRa+ cells, our scRNAseq map included predominantly epithelial cells and immune cells, most likely obtained due to the attachment of PDGFRa+ telopode fragments during the dissociation processes. Importantly, our extraction also yielded 329 pure mesenchymal cells (Fig. 4a-c, Methods).

**Fig. 4.**
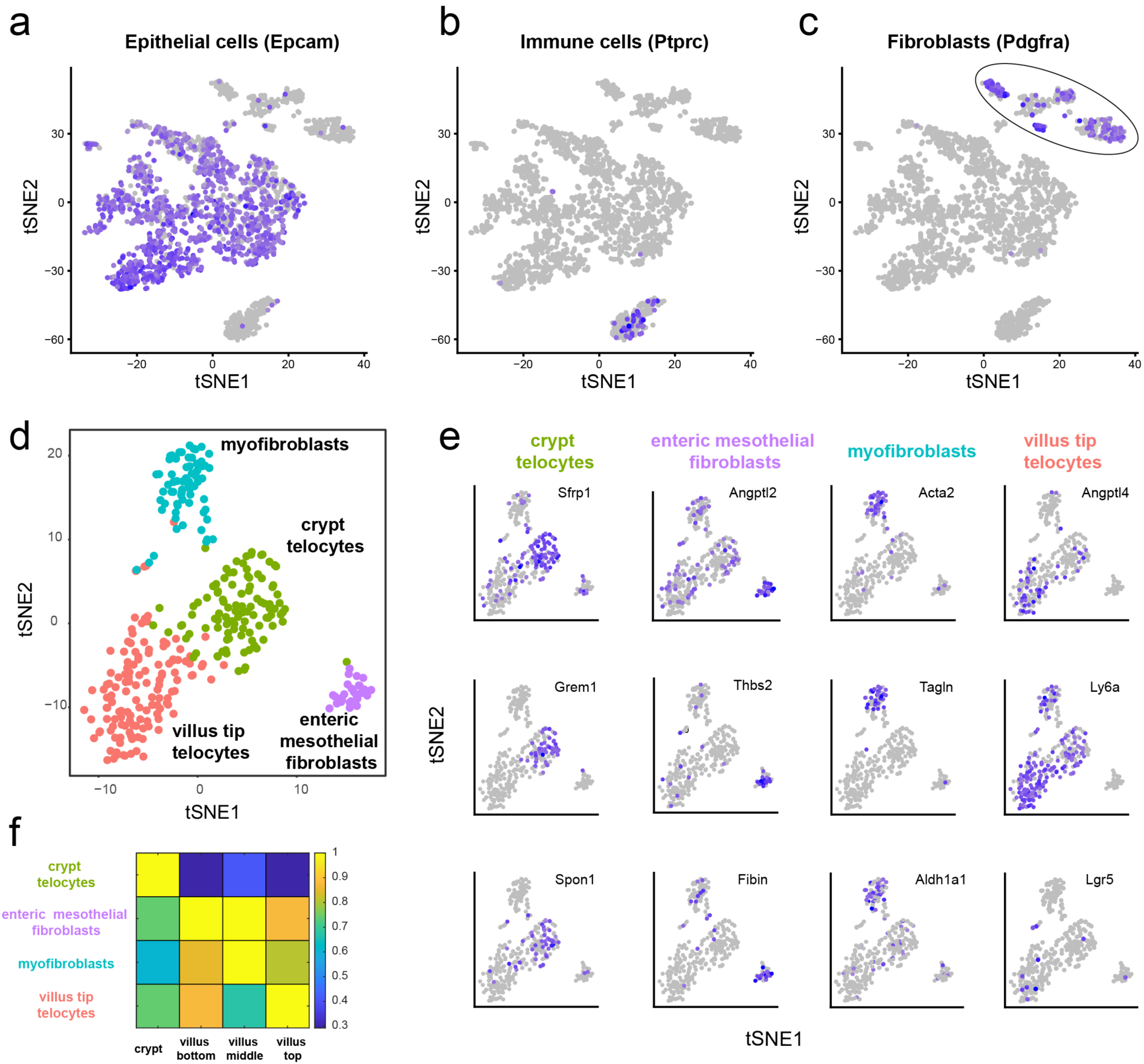
Single cell transcriptomics of the intestinal stroma identifies four spatially-stratified mesenchymal cell populations. a-c) tSNE plots of sequenced single cells highlighting the expression of Epcam, an epithelial marker (a), Ptprc, encoding CD45, a pan-immune marker (b) and Pdgfra (c) a mesenchymal marker. Purple hue indicates log expression levels. d) tSNE plot of the Pdgfra+ cells from the cluster circled in c) Colors denote the identified four clusters, obtained by re-clustering the Pdgfra+ cluster cells in c). e) tSNE plots colored by selected markers for the four mesenchymal cell clusters (see Extended Data Fig. 3 and Table S4 for the complete list of markers). Lgr5 is highly expressed in the Villus Tip Telocytes (7 out of 10 Lgr5+ cells belong to the VTT cluster, p=0.0121). f) Summed expression of the top markers for each mesenchymal cell cluster in the LCMseq data (Methods).

We focused on the mesenchymal cell clusters that were positive for PDGFRa. Reclustering these 329 cells yielded four distinct mesenchymal cell populations (Fig. 4d-e, Table S3). We used Seurat^29^ to identify distinct gene expression markers for each of the four cell populations (Fig. 4e, Extended Data Fig. 3, Table S4). To identify potential zonation of these mesenchymal cell types we used our LCMseq map (Table S1) to examine the expression of the marker genes from each cluster (Fig. 4f, Methods). Using the marker gene identities and their spatial patterns, we annotated the clusters as crypt telocytes, enteric mesothelial fibroblasts, myofibroblasts and villus tip telocytes (VTTs). 10 cells out of the 329 mesenchymal cells were Lgr5+, 7 of which were in the VTT cell population (Fig. 4e, hypergeometric p=0.0121). The remaining 3 Lgr5+ cells in our scRNAseq data were interspersed with 1 cell in each of the remaining three cell populations.

We used smFISH to validate the spatial patterns of expression of the cell type markers revealed by the scRNAseq measurements. Grem1 and Sfrp1 expression was elevated in telocytes at the bottom of the crypts (Extended Data Fig. 4a,c). Ly6a and Angptl4 expression was elevated in VTTs (Extended Data Fig. 4b,c). Notably, although the scRNAseq data suggested higher levels of Bmp2 and Bmp4 ligands in crypt telocytes, our smFISH measurements showed significantly higher levels of Bmp2 and Bmp4 in VTTs, in line with the LCM measurements (Fig. 1c, Extended Data Fig. 4c). Enteric mesothelial fibroblasts^27^ were marked by Thbs2, Fibin and Rgs5, as well as Angptl2, a gene previously shown to inhibit Bmp^30^. These were interspersed throughout the crypt-villus axis and were localized at the core of the villus, away from the epithelial layer (Extended Data Fig. 4d), as were myofibroblasts, marked by Acta2, Tagln and Aldh1a1 (Fig. 4e).

VTTs, as well as crypt telocytes, expressed high levels of ECM components such as Col3a1, Col1a1 and Timp2 (Table S3). VTTs expressed elevated levels of genes encoding microfibrillar proteins such as Mfap5, Emilin2 and Fbn1 (Fig. 5a, Extended Data Fig. 3, Table S3). Differential gene expression between crypt telocytes and VTTs further revealed elevated levels of the non-canonical Wnt ligand Wnt5a in VTTs (Fig. 5a), suggesting that VTTs implement a switch from canonical to non-canonical Wnt signaling^31,32^. Indeed, we observed broad expression of Axin2, a transcriptional target of canonical Wnt signaling, along the villi epithelial cells, with a decrease at the villus tip (Fig. 5c), the zone of stromal Wnt5a expression.

**Fig. 5.**
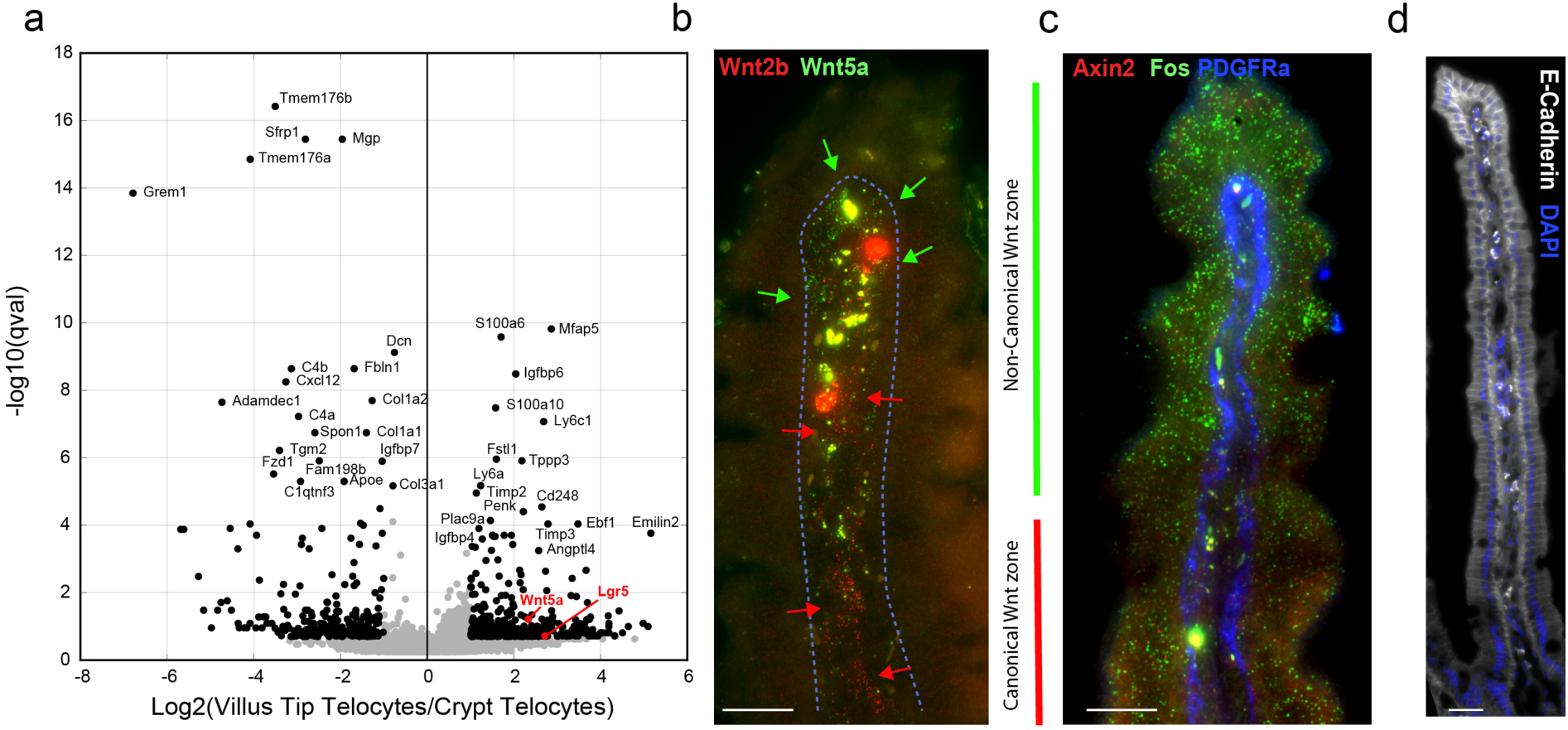
VTTs implement a spatial switch from canonical to non-canonical Wnt signaling. a) Volcano plot demonstrating differential gene expression between VTTs and crypt telocytes. Black dots have qval<0.2 and expression fold change larger than 2 or smaller than ½. Wnt5a and Lgr5 are marked in red. b) Spatial shift in the stroma from the expression of canonical Wnt2b (red dots, marked by red arrows) to non-canonical Wnt5a (green dots, marked by green arrows). Scale bar – 20 µm. c) Axin2 (red dots) is expressed broadly along the villus axis and repressed at the villus tip. Fos (green dots) is highly expressed in villus tip enterocytes. scale bar – 20 µm. d) E-Cadherin protein (gray), encoded by Cdh1 gene, is induced in villus tip enterocytes. Large blobs in B-D are autofluorescent signals originating in immune cells.

### VTT ablation perturbs epithelial gene expression at the villus tip

Enterocytes undergo profound gene expression changes as they approach the villus tip, a few hours before they are shed off into the lumen^4^. The villus tip enterocyte expression program includes induction of the adherens junction protein E-cadherin (Fig. 5d), the AP-1 transcription factor components Fos (Fig. 5c) and Jun, the transcription factor Klf4 and the immune-modulatory purine-metabolism program consisting of Ada, Nt5e and Slc28a2^4^. To assess the role of VTTs in regulating enterocyte zonation at the tip, we ablated Lgr5 cells using the Lgr5–GFP-DTR mouse model^25^, in which Lgr5+ cells express diphtheria toxin (DT) receptor and GFP. In this mouse model, 100% of Lgr5+ VTTs were positive for GFP (60 out of 60 cells counted over 10 villi from 2 mice, Fig. 3a,b, Fig. 6a, Extended Data Fig. 5a). 24 hours after DT administration, Lgr5+ VTTs were completely ablated, as were the crypt Lgr5+ stem cells (Fig. 6b). As previously shown, Lgr5 expression re-appeared in the crypt bottom 48 hours after ablation^25^, yet not at the villus tip, as evident by the complete loss of GFP signal (Fig. 6c) and Lgr5 mRNA (Extended Data Fig. 5).

**Fig. 6.**
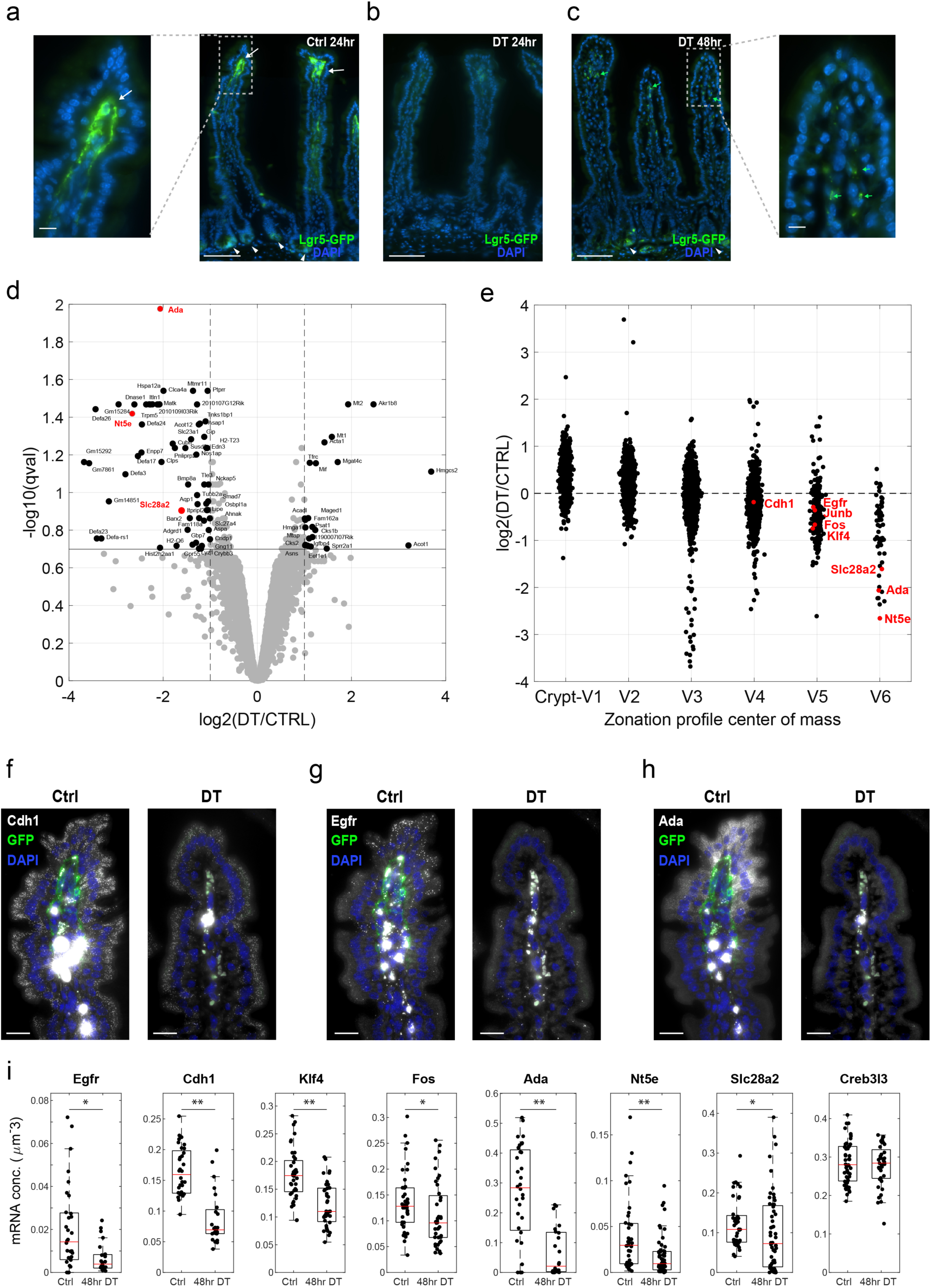
VTT ablation perturbs enterocyte expression at the villus tip. a) GFP fluorescence (green) observed in both crypt base columnar cells (CBCs, white arrowheads) and VTTs in Lgr5–GFP-DTR mice. Inset on left shows a blow up of villus tip with GFP+ VTTs (white arrow). b) Both VTTs and CBCs are ablated 24hr after DT administration, as evident by the lack of GFP fluorescence. c) GFP fluorescence re-appears in the crypt (white arrowheads) but not at the villus tip stroma 48hr after DT administration, indicating stable loss of VTTs. Inset on right shows a blow up of villus tip with no GFP+ VTTs. Green blobs are autofluorescent elements, also marked by green arrows. Scale bar in a-c – 100 µm, insets scale bar – 10 µm. d) Volcano plot demonstrating the changes in enterocyte gene expression 48hr following VTT ablation. Enterocyte villus tip genes that are reduced include Ada, Nt5e and Slc28a2 (red), composing the purine metabolism immune-modullatory tip module^4^. Black genes have q-values lower than 0.2 and max expression higher than 5*10^−6^ (Methods). e) Enterocyte genes normally induced at the villus tip are reduced in expression 48hr following VTT ablation. Correlation between change in expression and expression zone - R=-0.53, p<10^−60^. Enterocytes were classified into six villus zones as in Moor et. al.^4^. f-h) smFISH validations of enterocyte villus tip genes that are changed 48hr following VTT ablation – in each panel Ctrl-left, DT-right. f) Cdh1. g) Egfr. h) Ada. Scale bar in f-i – 20 µm i) Quantification of smFISH experiments, demonstrating that key epithelial villus tip genes are repressed 48hr after VTT ablation (Egfr, Cdh1, Klf4, Fos, Ada, Nt5e, Slc28a2 **p<10^−3^ *p<0.05), whereas others remain unchanged (Creb3l3, p=0.5). Analysis was performed on two mice. Boxes show 25-75 percentiles of the smFISH expression, horizontal red lines are medians.

Ablation of Lgr5+ crypt base columnar cells has been shown to initiate a regenerative program in the remaining crypt cells^33^. Importantly, migration of epithelial cells from the crypt to the villus tip takes 3-5 days^1,3^. We therefore argued that at short times following ablation of Lgr5+ cells, reduction in the expression of genes normally confined to the villus tip epithelium would be a result of loss of signals from the Lgr5+ VTTs, rather than incoming enterocytes, the state of which was perturbed in the crypt. To obtain a global view of the impact of VTTs on villus tip enterocyte gene expression, we therefore performed RNA sequencing of the epithelial layer 48 hours after DT administration (Table S5, Fig. 6d,e, Methods). We identified a significant repression in enterocyte genes that were normally zonated towards the tip (Spearman correlation of R=-0.53, p<10^−60^ 48hr post ablation between expression change following ablation and expression zone, Fig. 6e, Table S6). Villus tip enterocyte genes that were strongly reduced in expression upon VTT ablation included the purine metabolism module genes Slc28a2, Nt5e and Ada (Fig. 6d-i).

Additional prominent villus tip enterocyte genes include the transcription factors Klf4 and Fos, Cdh1, encoding E-cadherin and Egfr^4^. These genes have a relatively high basal expression level either throughout the villi axis (Cdh1, Klf4 and Fos) or at the crypt (Egfr). Expression changes in enterocyte at the villus tip could thus be masked for such genes in bulk RNA sequencing (Fig. 6d). To assess their expression changes, we therefore performed smFISH and measured expression specifically at the tip enterocytes, identifying a significant reduction of Egfr, Cdh1, Klf4 and Fos, in addition to Ada, Nt5e and Slc28a2 (Fig. 6g-i). Creb3l3, an enterocyte gene that is elevated at the villi tip, did not exhibit changes in expression upon VTT ablation. Epithelial Axin2 expression did not change following VTT ablation (Table S5). These results demonstrate that VTTs are important regulators of the spatial expression programs of enterocytes at the villus tip, instructing the epithelial expression of key genes such as Egfr, Cdh1, Klf4, Fos, Nt5e, Ada and Slc28a2.

To assess the long-term consequences of VTT ablation we examined small intestinal tissue three weeks after VTT ablation. At this time point, VTTs re-appeared in 65% of the villi tips (Fig. 7, 46 GFP+ villi out of 71 villi, counted over 2 mice). Notably, the mRNA levels of the enterocyte villus tip genes Ada, Nt5e, Egfr, Fos and Klf4 remained significantly lower in the villi tips that lacked VTTs, compared to tips where VTTs reappeared (Fig. 7, mRNA conc. more than 2-fold higher in GFP+ vs. GFP-villi). In contrast, Cdh1 and Creb3l3 exhibited only a slight, yet statistically significant higher expression in the GFP+ villi (1.2 fold and 1.13 fold for Cdh1 and Creb3l3 respectively in GFP+ vs. GFP-). The correlation between VTT re-appearance and villus tip epithelial gene expression at this time point, when crypts have already returned to normal homeostatic state, further supports the instructive role of VTTs in regulating the enterocyte villus tip expression program.

**Fig. 7.**
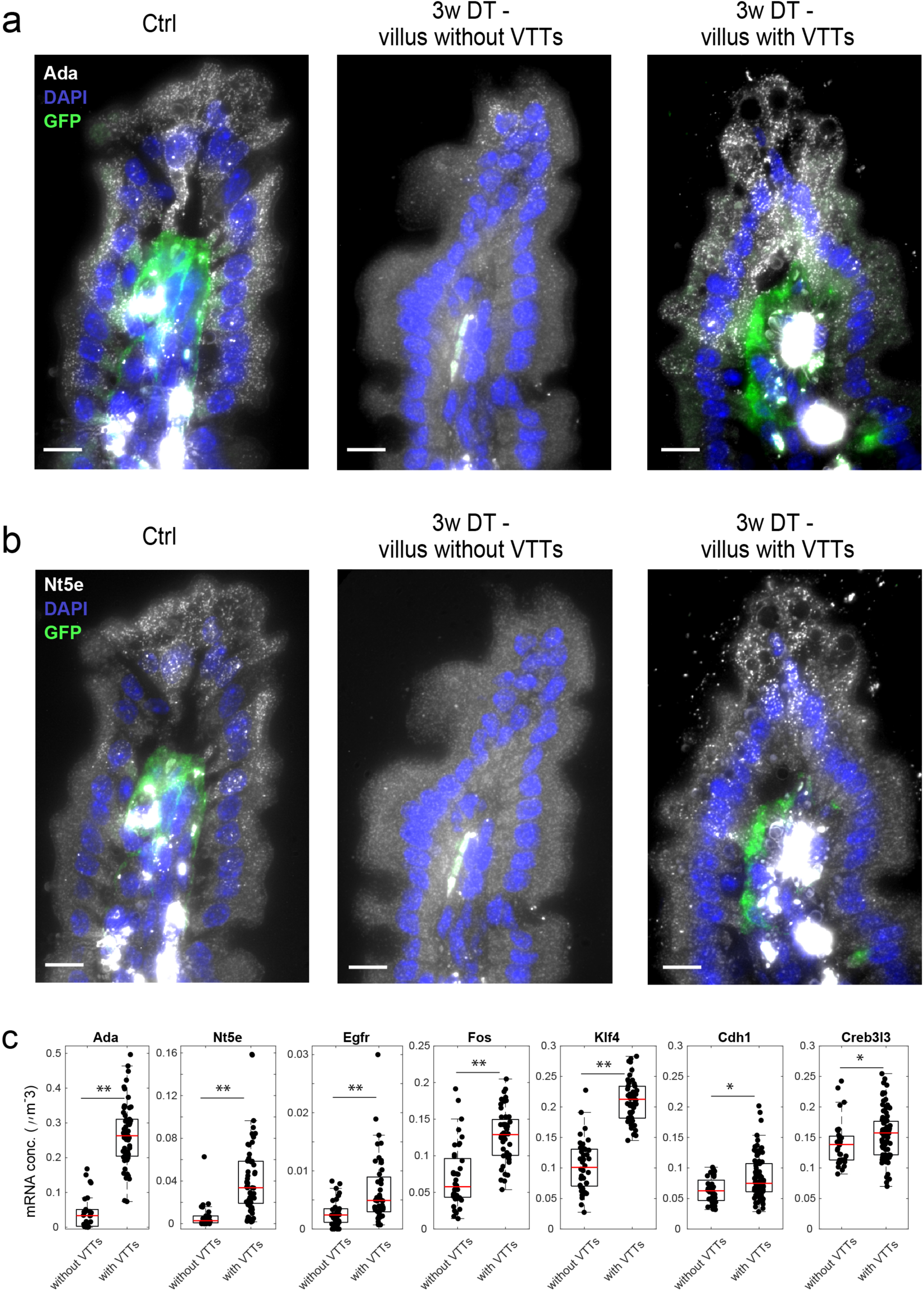
Analysis of the villus tip epithelial cells of Lgr5–GFP-DTR 3 weeks after VTT ablation shows correlation between the re-appearance of VTTs and the expression of enterocyte tip genes. a) smFISH example of Ada showing higher expression in the villus tip with VTTs compared to villus tip without VTTs after 3 weeks. Scale bar 10 µm. b) smFISH example of Nt5e showing higher expression in the villus tip with VTTs compared to villus tip without VTTs after 3 weeks. Scale bar 10 µm. c) Quantification of expression differences between villi with VTTs and villi without VTTs. Measurements were performed over 15 villi per mouse for two mice. (*p <0.05, **p <0.001).

## Discussion

Our study exposed the spatial diversity of mesenchymal cells along the crypt-villus axis. We identified a cell population of Lgr5+ VTTs that form a highly localized niche for the villus tip epithelium, facilitating the massive transcriptional changes that enterocytes undergo before they are shed from the tissue^4^. VTTs are a source of Bmp ligands (Fig. 2d,e, Extended Data Fig. 4c, Table S1) and may implement a switch from canonical to non-canonical Wnt signaling, through the expression of the non-canonical Wnt ligand Wnt5a (Fig. 5b). Similar spatially antagonistic expression of stromal Wnt5a expression and canonical Wnt activity has been recently demonstrated in the mouse prostate^31^. This switch may be important for the induction of E-cadherin, a major component of the adherens junction that is strongly induced at the villus tip enterocytes (Fig. 5d)^34^. Wnt5a is also an activator of the AP1 transcription factors^35^, composed of the enterocyte villus tip genes Fos and Jun^4^. Future work will resolve the precise molecular mechanisms involved in the induction of specific enterocyte villus tip genes by VTTs.

VTTs are ideally suited to orchestrate gene expression programs along the intestinal villi. Their close proximity to the epithelial sheet and their intricate extensions towards the lamina propria facilitate close interactions with both epithelial and other stromal cell types. Unlike the short-lived epithelial cells and the often mobile immune cells, VTTs are static, quiescent and long-lived, and can therefore integrate long-term information related to diets and pathologies. The intestine exhibits a remarkable ability to adapt to such perturbations^36,37^. Future work will resolve the roles of VTTs in shaping villus epithelial programs in such perturbed states. The re-appearance of VTTs 3 weeks following their ablation (Fig. 7) supports their importance for preserving the normal epithelial function. It will be important to identify the cellular origins of these re-appearing VTTs. Our study exposed VTTs as major regulators of the expression of villus tip enterocyte genes (Fig. 6e), yet enterocyte gene expression changes throughout the villus axis^4^. It will be interesting to explore the additional effects of zonated luminal signals and potential enterocyte ‘internal clocks’ in shaping enterocyte zonation.

Lgr5+ fibroblasts have previously been identified in the lung^38^, suggesting that Lgr5+ stromal cells may constitute a localized mesenchymal niche component in other tissues. It will be important to use similar spatial transcriptomics approaches to explore the cross-talk between the mesenchymal cell populations and zonated immune cell populations^26^ as well as other stromal components such as enteric neurons^27^. Our finding of positive expression of the Lgr5-EGFP-IRES-creERT2 knock-in as well as the Lgr5–GFP-DTR knock-in cassettes at the villus tip is important for interpreting physiological effects of perturbations driven by the Lgr5 promoter, such as lineage tracing, cell ablation and conditional knock-outs.

Recent efforts for establishing tissue cell atlases rely on the ability to deconvolve the complex spatial interactions of distinct parenchymal and non-parenchymal cell types in each tissue^13^. Our study performs such combined analysis in the mouse small intestine, a prototypical zonated tissue. Similar approaches could be applied to reveal zonated interactions between stroma and epithelial cells in other structured organs.

## Extended Data Figures

**Extended Data Figure 1.**
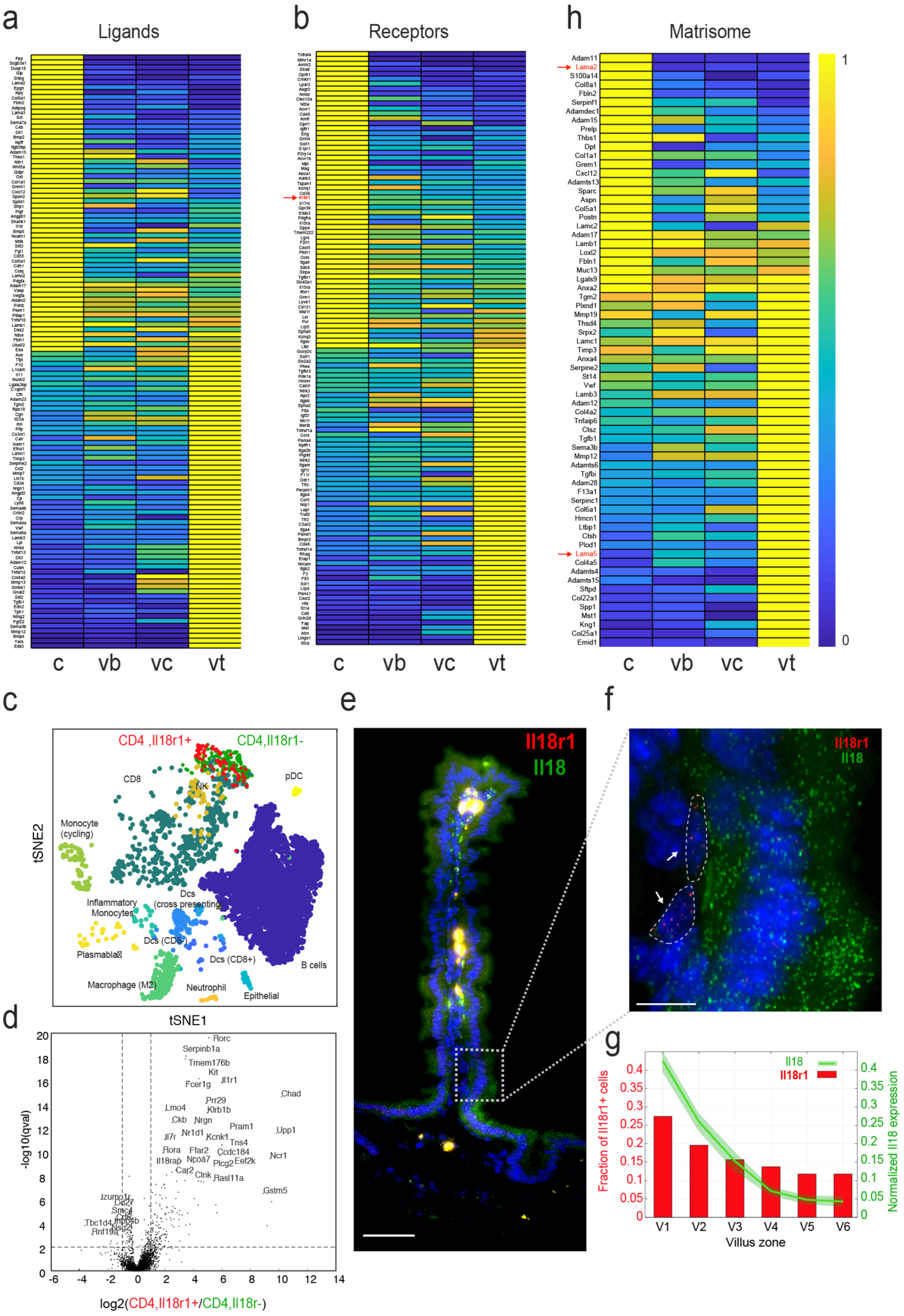
Zonated expression of stromal ligands (a) and receptors (b). Il18r1 marked in red. c) tSNE plot of immune cells in the small intestine taken from Biton et. al.^26^. Il18r1 is expressed in a subset of a cluster annotated as CD4 (red cells). d) Differential gene expression between the Il18r1+ cells (red cells in c)) and Il18r1-cells of the same cluster (green cells in c)). These genes are markers of innate lymphoid cells type 3. The y axis was truncated at qval=10^−20^, leaving out Il18r1 (having expression ratio of 4,240, qval=10^−79^). Vertical lines denote log2 ratios of 1 and −1. Horizontal line denotes qvalue of 0.01. e) smFISH for Il18 (green) and Il18r1 (red) demonstrating spatial colocalization at the lower villus zones. Scale bar – 50µm. f) blow-up of a region boxed in e) demonstrating spatial adjacency of stromal immune cells with Il18r1 transcripts (red dots, cells marked by red arrows and outlined in red) and Il18+ villus bottom epithelial cells. Scale bar – 10 µm. g) Quantification of the fraction of Il18r1+ cells (left y-axis) along the villus axis showing colocalization to the region where Il18 is highly expressed (right y-axis). Il18 expression is taken from Moor et. al.^4^, normalized to sum to 1 over villus zones. h) Zonated expression of proteins constituting the extracellular matrix (ECM, also termed the ‘matrisome’^17^). Red arrows highlight the crypt-zonated Lama2 and villus tip-zonated Lama5, previously described to be inversely zonated^18,19^. Each row in (a,b,h) was scaled by its max expression across the four zones.

**Extended Data Figure 2.**
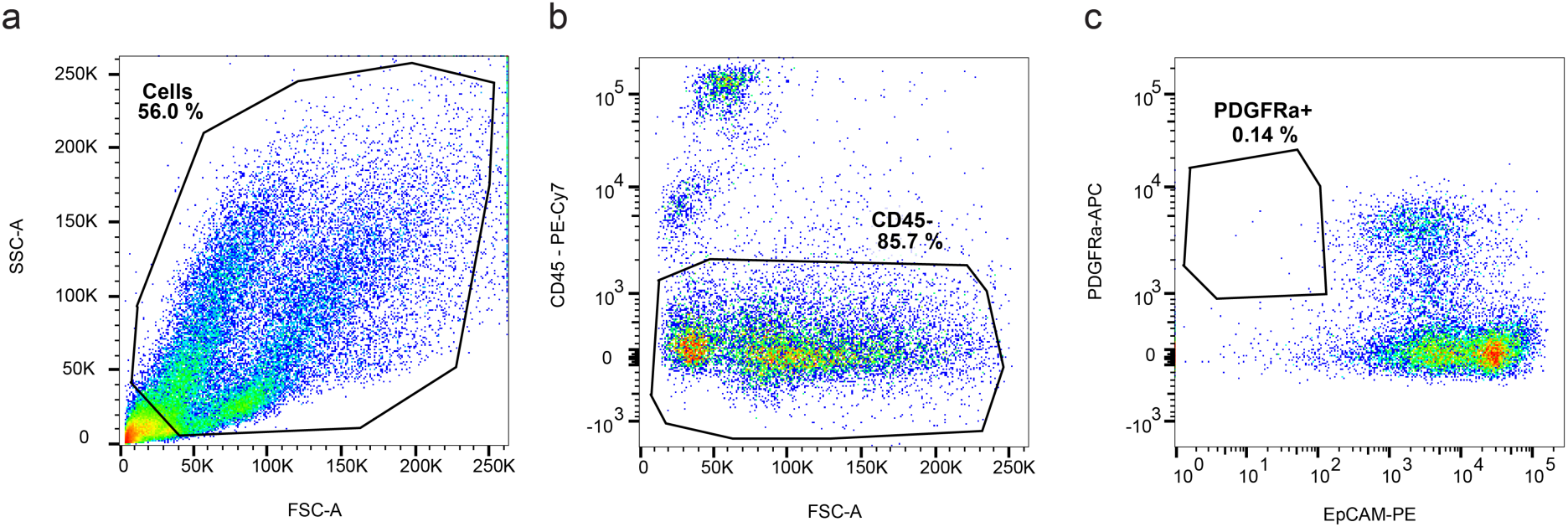
FACS gates used to enrich for PDGFRa+ cells that included the intestinal telocytes, as well as other mesenchymal cell types. a) FSC-A and SSC-A were used to select cells based on size. b) CD45 was used to gate out immune cells. c) PDGFRa and EpCAM fluorescense were used to select EpCAM negative, PDGFRa positive telocytes. Numbers in a-c) represent the percent of gated cells.

**Extended Data Figure 3.**
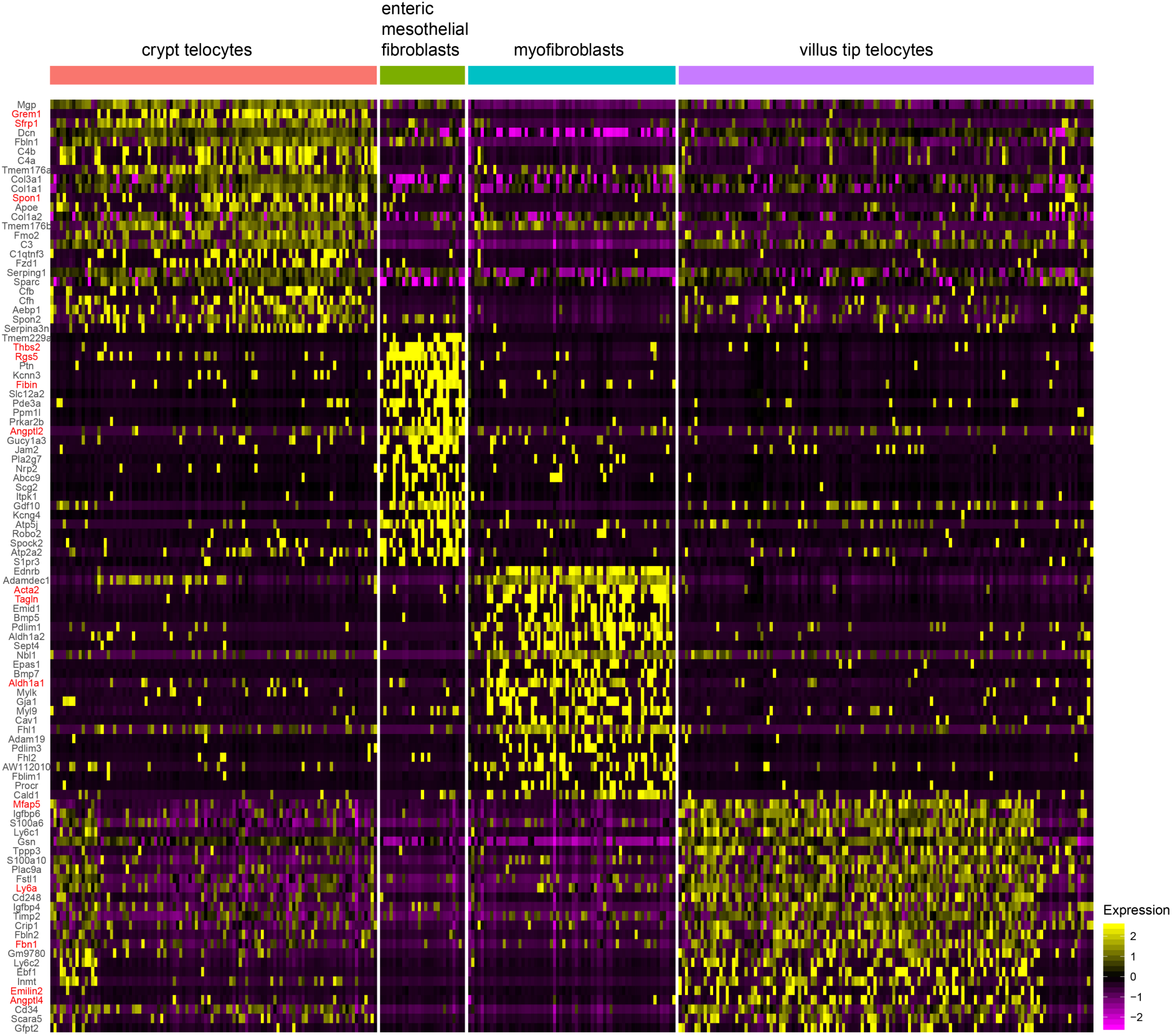
Heatmap showing the 25 top differentially expressed genes in each mesenchymal cell population - crypt telocytes, enteric mesothelial fibroblasts, myofibroblasts, villus tip telocytes. Genes discussed in the main text are marked in red.

**Extended Data Figure 4.**
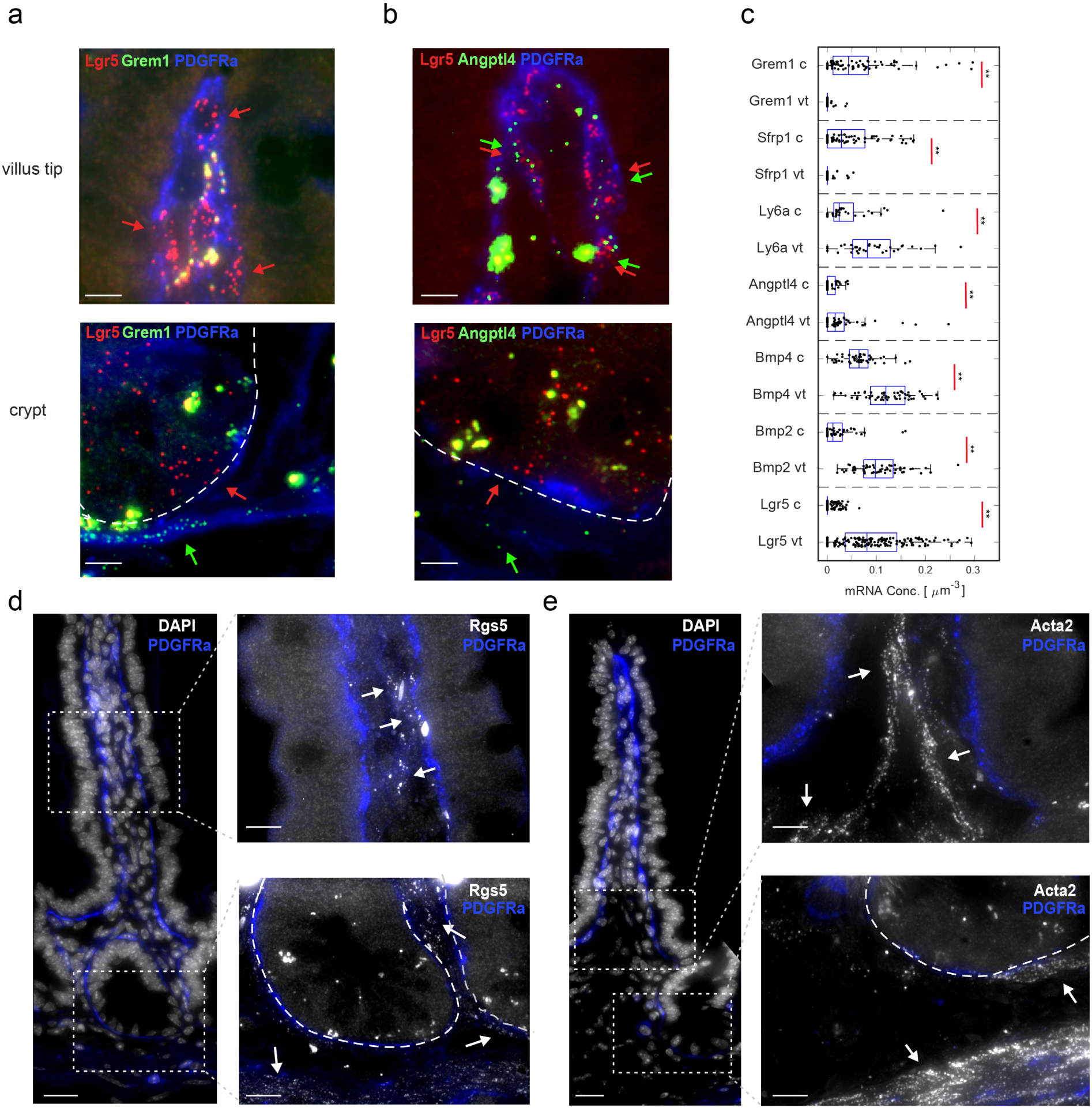
a) Grem1 mRNAs (green dots, green arrows mark positive cells) are expressed in crypt telocytes and not in VTTs. Scale bar – 5 µm. b) Angptl4 (green dots) are expressed more highly in VTTs compared to crypt telocytes. Scale bar – 5 µm. Red dots are Lgr5 mRNAs (red arrows mark positive cells), blue is PDGFRa antibody staining. c) Quantification of smFISH validations for crypt telocytes and VTT markers. ** denote pvalue<10^−4^ from two sided Wilcoxon ranksum test, c – crypt, vt – villus tip. Boxes show 25-75 percentiles of the smFISH expression, vertical lines are medians. d) Enteric mesothelial fibroblasts are scattered throughout the crypt-villus axis. Grey dots are mRNAs of Rgs5. blue is PDGFRa antibody staining. Scale bars – 20 µm, insets scale bar – 10 µm e) Myofibroblasts are scattered throughout the crypt-villus axis and localized away from the epithelial layer. Grey dots are mRNAs of Acta2. Blue is PDGFRa antibody staining. Scale bars – 20 µm, insets scale bar – 10 µm. White arrows mark positive cells in d) and e). White dashed line mark crypt borders.

**Extended Data Figure 5.**
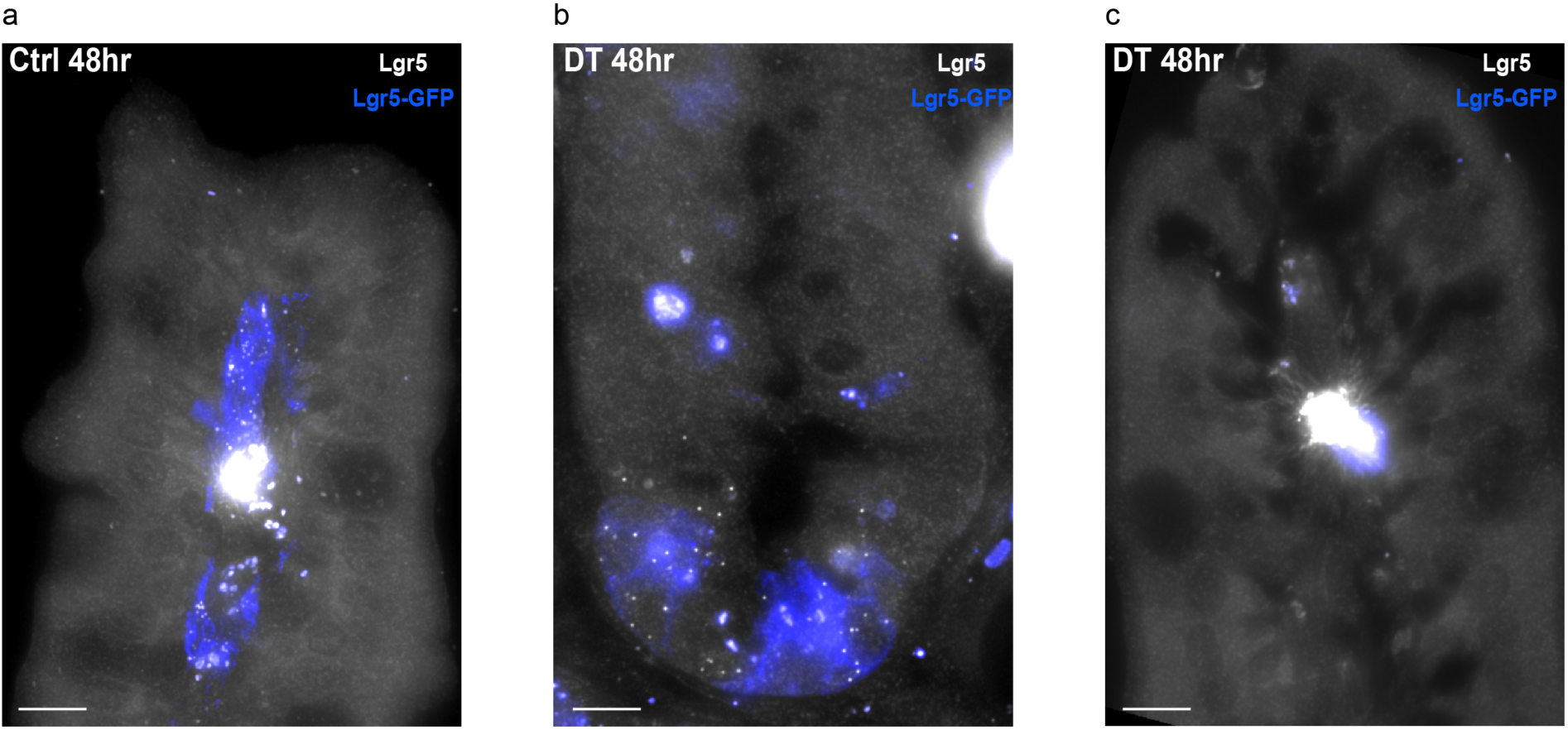
Villi tips are devoid of GFP and Lgr5 mRNA in Lgr5–GFP-DTR mice 48hr after VTT ablation. a) Bright dots are single mRNA of Lgr5 co-localized with GFP positive cells. b) GFP and Lgr5 reappear at the crypt stem cells. c) Villi tips are devoid of GFP and Lgr5 mRNA. Scale bar 10 µm.

## Supplementary Tables

Supplementary Table 1 – Average gene expression in the laser capture microdissected stromal zones. For each sample, UMI counts were normalized to the sum for all non-enterocyte genes that individually take up less than 1% of the total sample counts (Methods). TPM values presents the complete non-filtered data.

Supplementary Table 2 – Correlations between the stromal and epithelial spatial expression domains of ligands and their respective receptors. Epithelial expression was taken from Moor et al.^4^, Stromal expression taken from Table S1.

Supplementary Table 3 – Average expression of the four mesenchymal cell populations based on the scRNAseq data. Expression is in units of fraction of total UMIs per cell.

Supplementary Table 4 – Markers for the four mesenchymal cell popluations obtained from Seurat^29^.

Supplementary Table 5 – UMI counts for bulk RNAseq of epithelial cells extracted from DT-treated and mock-treated Lgr5–GFP-DTR mice 48 hrs after ablation.

Supplementary Table 6 – Differential gene expression between epithelial cells extracted from DT-treated and mock-treated Lgr5–GFP-DTR mice 48 hrs after ablation.

Supplementary Table 7 – Sequences of the smFISH probes libraries used in this study.

## Methods

### Mice and tissues

All animal studies were approved by the Institutional Animal Care and Use Committee of WIS and UCSF. C57bl6 male mice age 8-10 weeks were obtained from the Harlan laboratories, all mice were fed regular chow ad libitum. Mice were sacrificed by cervical dislocation. For LCM, jejunum tissues were harvested and embedded directly in OCT without fixation. For smFISH, jejunum tissues were harvested and fixed in 4% formaldehyde for 3 hr, incubated overnight with 30% sucrose in 4% formaldehyde and then embedded in OCT in the form of swiss-rolls. 7 µm cryosections were used for hybridization. Mouse jejunum cells for single-cell RNAseq were extracted from eight mice. All smFISH quantifications were performed on at least 2 mice. Images in Figure 3C,D were taken from jejunum tissues of Lgr5-EGFP-IRES-CreERT2 mice^2^. *Lgr5*^*DTRGFP*^ (Lgr5-GFP-DTR) mice^25^ were housed in the University of California San Francisco (UCSF) animal facility in compliance with all ethical guidelines established by the Institutional Animal Care and Use Committee (IACUC) and Laboratory Animal Resource Center.

### Antibodies used in this study

The following antibodies were used for FACS cell isolation: CD31 (PE-Cy7 102418, Biolegend), CD45 (APC-Cy7 103116, PE-Cy7 103114, Biolegend), PDGFRa (Alexa 488 FAB1062G, R&D systems, APC 135908, Biolegend), Epcam (PE 118205, Biolegend). All FACS antibodies were used at 1:100 dilution in the presence of TruStain fcX blocker (101320, Biolegend, 1:50). For PDGFRa immunofluorescence staining, goat anti mouse PDGFRa was used as first antibody (AF1062 R&D systems) at a concentration of 8 µg/µl in the smFISH hybridization buffer, Alexa fluor 488 conjugated Donkey anti goat (705-545-147 Jackson laboratories, 1:400) was used as secondary antibody. For E-cadherin immunofluorescence staining FITC anti E-Cadherin (612131 BD Biosciences) was used at dilution of 1:100.

### Laser capture microdissection (LCM)

Tissue blocks for laser-capture microdissection were freshly prepared at the morning of the collection day. The jejunum part was transected on top of Wattman paper soaked in PBS to support the opened structure in petri dish on ice. The tissue was washed with cold PBS and embedded in OCT on dry ice. LCM protocol was applied as previously described^4,39^ with minor modifications. Sections of 9µm thickness were cut from the frozen block, mounted on polyethylene-naphthalate membrane-coated 518 glass slides (Zeiss, 415190-9081-000), air-dried for 20sec at room temperature, washed in 70% ethanol for 25sec, incubated in water for 25sec (Sigma-Aldrich, W4502), stained with HistoGene Staining Solution for 20sec (ThermoFisher Scientific, KIT0401), washed again in water for a of 25sec. The stained sections were dehydrated with subsequent 25sec incubations in 70%, 95% and 100% EtOH and air-dried for 60s before microdissection. Tissue sections were examined using a bright field imaging microscope (Observer.Z1, Zeiss), and microdissected with a UV laser-based unit (PALM-Microbeam, Zeiss). To ensure minimal damage to the surrounding cells, laser intensity and focus were calibrated before each session using Zeiss calibration wizard supplemented with the LCM operating software (Zeiss). Manual detection of analyzed regions in each tested slide and labeling of the desired areas was done with PALM 10X and 20X lenses. Tissue fragments were catapulted and collected in 0.2ml adhesive cap tubes (Zeiss, 415190-9191-000). Each capture section was visually confirmed by focusing the PALM on the targeted adhesive cap after the collection session and immediately resuspend in 10.5µl of 1x reaction buffer from SMART-Seq v4 Ultra Low Input RNA Kit (Clontech, 634888). Tissue lysis was achieved by incubation for 5min at room temperature and stored at −20°C. Collected samples were stored at −80°C until the library preparation. Three distinct zones spanning approximately equal heights from the crypt-villus border to the villus tip, were collected from long villi (about 500µm) - villus bottom, villus center and villus top (Fig. 1). An area of ∼30,000 μm^2^ per zone was collected for each mouse by pooling 9-11 distinct villi from a single tissue section. For mice 1-3 (Table S1) we additionally catapulted crypt stroma and extracted material from two sequential tissue sections for each zone, with the exception of the villus center zone of mouse 3, where one section failed to amplify (Table S1, TPM value tab).

### SMART-Seq for bulk LCM samples

RNA libraries from the bulk tissues were prepared using SMART-Seq v4 Ultra Low Input RNA Kit (Clontech, 634888), with a 15 PCR cycles for the amplification step. Subsequent steps were applied as mentioned in the protocol. Nextera XT DNA Library Prep Kit (Illumina) was use to finalize the libraries. Library concentration and quality control were determined using NEBNext Library Quant Kit (New England Biolabs) and Agilent High Sensitivity D1000 ScreenTape System (Agilent, 5067-5584). Library final concentration of 2.4pM was loaded on NextSeq550 (Illumina) sequencing machine aiming for 20M reads per sample with the following cycle distribution: 38bp read1, 8bp index1, 8bp index2, 38bp read2.

### Stroma LCM bulk RNA-seq analysis

Illumina raw files were converted to FASTQ files using bcl2fastq 2.17 (Illumina), following by pseudoalignment with Kallisto 0.43.0^40^ to a transcriptome index of the GRCm38.94 (Ensembl), filtered to transcripts with “coding_genes” entry. The following flag was used for kallisto: --b 100 (the number of bootstrap samples), --rf-stranded (strand specific reads, first read reverse). Sleuth 0.30.0^41^ running on R 3.5.1 was utilized to create a TPM table (Transcripts Per Million) for each sample (Table S1).

### Stroma LCM data processing

We removed 31 enterocyte genes, determined as genes the summed fraction of which makes up the top 50% of enterocyte expression. To this end we considered the maximal expression of all genes in enterocytes over the zones as measured in Moor et al.^4^. We normalized the TPM values upon removing these genes by the sum of all remaining TPM. We next divided the gene expression values in each sample by the sum for all genes that individually make up less than 1% of the sample’s summed expression values. For each of the four zones, we computed the means and standard errors of the means over the different samples. Ligand receptor pairs were extracted from Ramilowsky et al.^12^, Matrisome components were extracted from Naba et al.^17^. Only genes with maximal zonation value larger than 5*10^−6^ were considered.

### Analysis of Il18r1 stromal cells

We used the single cell RNAseq data of Biton et al.^26^ (6558 cells annotated as WT control, Extended Data Fig. 1c). Il18r1 positive cells belonged to a cluster annotated as ‘CD4’. We performed differential gene expression using Wilcoxon rank-sum tests between the Il18r1+ and Il18r-cells in that cluster (Extended Data Fig. 1d). We used Immgen My Geneset tab (https://www.immgen.org) to annotate the 200 genes that had the highest fold-change between Il18r1+ cells and Il18r1-cells within the CD4-annotated cluster (only genes with expression above 5*10^−5^ were used for this analysis). The gene set was enriched in innate lymphoid cells type 3.

### Hybridization and imaging

Probe libraries were designed using the Stellaris FISH Probe Designer Software (Biosearch Technologies, Inc., Petaluma, CA). 7µm thick sections of fixed Jejunum were sectioned onto poly L-lysine coated coverslips and used for smFISH staining. The intestinal sections were hybridized with smFISH probe sets according to a previously published protocol^15^. SmFISH probe libraries (Table S7) were coupled to Cy5, TMR or Alexa594. Telocytes cells were detected by PDGFRα primary antibody that was added to the smFISH hybridization buffer and Alexa fluor 488 conjugated Donkey anti goat as secondary antibody in GLOX buffer for 20 minutes after DAPI (Sigma-Aldrich, D9542) nuclear staining. All images were taken as scans extending from villus tip to crypt bottom using 100x magnifications, hence several fields of view were stitched together to cover the whole crypt-villus unit. Stitching was performed with the fusion mode linear blending and default settings of the pairwise stitching plugin^42^ in Fiji^43^. Quantification of smFISH was done using ImageM^44^. For smFISH quantifications, results were based on at least 30 cells from each region and from at least 2 mice. Dots were counted in the first 3μm of the Z-stack, and divided by the segmented cell volume to obtain the mRNA concentration per cell. Two-sided Wilcoxon rank-sum tests were used to assess significance. Quantifications of the proportions of Lgr5+ VTTs positive for the EGFP knock-in were done on jejunum tissues of Lgr5-EGFP-Ires-CreERT2 mice ^2^.

### Single cell isolation

Due to the difficulties of isolating intestinal stromal cells we applied several dissociation protocols and surface marker staining to obtain a broad sampling of cells. For all mice, cells were isolated from the jejunum. The jejunum was extracted and rinsed in cold PBS. The tissue was opened longitudinally and sliced into small fragments roughly 2 cm long and incubated in 10mM EDTA-PBS on ice for 10 min. The tissue was then moved to warm 5ml 10mM EDTA-PBS containing Liberase TM (100 μg/mL, Sigma) and DNaseI (2 U/mL, Sigma) and incubated at 37°C for 20 min while shaken vigorously every few minutes. At the end of the incubation time, 5ml of cold PBS was added to the cell suspension. The supernatant was filtered through a 100μm filter and centrifuged at 300 g for 5 min. the pellet was resuspended in FACS buffer (2mM EDTA, 0.5% BSA in PBS) and stained with the required antibodies for flow cytometry sorting.

Isolation of telocytes was performed as previously decribed in^9^. Briefly, Jejunum were dissected and washed thoroughly with Hank’s balanced salt solution (HBSS) and were incubated in 5 mM EDTA in HBSS for 10 min at 4 °C. Intestinal villi were scraped off using a coverslip and the remaining tissue was cut into small pieces and incubated in 5 mM EDTA and HBSS on ice for 10 min while pipetting to completely remove the remaining epithelium. After vigorous washes, the remaining mesenchymal fraction was incubated with 6 mg/ml Dispase II/0.05% trypsin solution (Sigma-Aldrich, 04942078001) supplemented with 1 U/ml DNaseI (Sigma) at 37 °C, until the solution became cloudy and the mesenchyme was dissociated (8 min). At the end of the incubation time, 3ml of warm FCS were added to the cell suspension to stop the digestion. Supernatant was filtered through a 70-μm strainer, centrifuged at 3500 rpm for 5 min. the pellet was resuspended in FACS buffer (2mM EDTA, 0.5% BSA in PBS) and stained with the required antibodies for flow cytometry sorting.

### Single-cell sorting

Single cells were sorted with SORP-FACSAriaII machine (BD Biosciences) using a 100 μm nozzle. Dead cells were excluded on the basis of 500 ng/ml Dapi incorporation. Sorte cells were negative for CD45 and EpCAM and positive for PDGFRa. Cells were sorted into 384-well cell capture plates containing 2 μl of lysis solution and barcoded poly(T) reverse-transcription (RT) primers for single-cell RNA-seq^28^. Barcoded single cell capture plates were prepared with a Bravo automated liquid handling platform (Agilent) as described previously^28^. Four empty wells were kept in each 384-well plate as a no-cell control during data analysis. Immediately after sorting, each plate was spun down to ensure cell immersion into the lysis solution, snap frozen on dry ice and stored at −80°C until processed.

### Massively Parallel Single Cell RNA-Seq (MARS-Seq) library preparation

Single cell libraries were prepared, as described in^28^. Briefly, mRNA from cells sorted into MARS-Seq capture plates were barcoded and converted into cDNA and pooled using an automated pipeline. The pooled sample was then linearly amplified by T7 in vitro transcription and the resulting RNA was fragmented and converted into sequencing ready library by tagging the samples with pool barcodes and Illumina i7 barcode sequences during ligation, reverse transcription and PCR. Each pool of cells was tested for library quality and concentration was assessed as described in^28^. Machine raw files were converted to fastq files using bcl2fastq package, to obtain the UMI counts, reads were aligned to the mouse reference genome (GRCm38.84) using zUMI packge ^45^ with the following flags that fit the barcode length and the library strandedness: -c 1-7, -m 8-15, -l 66, -B 1, -s 1, -p 16. This analysis resulted in 2,217 cells.

### scRNAseq data processing

For each single cell and for each gene we first subtracted the estimated background expression. Background was calculated for each 384-well plate separately, as the mean gene expression in the four empty wells. After subtraction, negative values were set to zero. Next, cells with total UMI counts lower than 400 or higher than 8,000 and total gene counts lower than 250 were removed. We used Seurat v2.3.4 package in R^29^ v3.5.3 to visualize and cluster the single cell RNAseq data (Fig. 4, Extended Data Fig. 3). Gene expression measurements (UMIs per gene) were normalized for each cell by the summed UMI, multiplied by a scale factor 10,000, and then log-transformed. To avoid undesired sources of variation in gene expression, we used Seurat to regress out cell-cell variation driven by total number of UMIs, and mitochondrial genes fraction. For detection of variable genes, we set a bottom cutoff of 0.25 and a top cutoff of 4 on the regressed log-transformed average gene expression, as well as a bottom cutoff of 0.5 on the dispersion. Cell clustering was based on PCA dimensionality reduction using the first 11 PCs, and a resolution value of 2.

We used cell type-specific markers to interpret the resulting 3 main clusters, Epcam was highly expressed in the epithelial cells clusters, Ptprc was highly expressed in the immune cells clusters and Pdgfra was highly expressed in the fibroblast cells clusters.

We next focused on the cell clusters that had an average expression of Pdgfra that was higher than 10^−4^ per cell (fraction of UMI counts). These included 329 cells. For this new set of cells, we re-ran Seurat with the following parameters; For detection of variable genes, we set a bottom cutoff of 0.25 and a top cutoff of 4 on the regressed log-transformed average gene expression, as well as a bottom cutoff of 1 on the dispersion. Cell clustering was based on PCA dimensionality reduction using the first 5 PCs, and a resolution value of 0.7. Marker genes were detected with Seurat FindAllMarkers function with parameters min.pct=0.2, logfc.threshold=0.25 (Table S4).

To identify zonation of the four mesenchymal cell types we examined the summed expression of concise sets of markers that were specific to each cell type. To select these markers, we included genes that had an average expression higher than 5*10^−6^ and were expressed in at least 10 cells in the respective cell type. We sorted these genes by the fold change of their expression levels compared to the maximal average expression in the other three mesenchymal cell types. We included the top 20 genes and further excluded genes for which the fold change ratio was lower than 5-fold. Differential gene expression between crypt telocytes and VTTs (Fig. 5a) considered all genes with average expression larger than 10^−5^ of cellular UMIs and more than 5 cells positive for the gene in at least one of the two populations. Storey’s method was used to compute q-values.

### Lgr5+ cell ablation

For Lgr5+ cell ablation, male *Lgr5*^*DTRGFP*^ mice aged 4-5 months were administered 50 μg/kg diphtheria toxin (322326 Sigma) or saline vehicle intraperitoneally. Sample were collected 24 or 48 hr post injection. For the 24 hr time points, four DT-injected and two mock-injected mice were used for histological examination (Fig. 6). For the 48 hr time point, four DT-injected and four mock-injected mice were used for histological examination, smFISH and epithelial cell sequencing. Additionally, two DT-injected and two mock-injected mice were sacrificed 20 days after administration and used for histological examination and smFISH. For epithelial cell isolation, 3 cm from the proximal jejunum were extracted and rinsed in cold PBS. The tissue was opened longitudinally and sliced into small fragments roughly 2 cm long and incubated in 10mM EDTA-PBS on ice for 20 min. The tissue was then moved to warm 5ml 10mM EDTA-PBS and incubated at 37°C for 5 min while shaken vigorously every few minutes. At the end of the incubation time, 5ml of cold PBS was added to the cell suspension. The supernatant was filtered through a 100μm filter and centrifuged at 300 g for 5 min. the pellet was snap frozen in liquid nitrogen and was processed for bulk RNA sequencing. For smFISH blocks, 5 cm from the distal jejunum was fixed immediately in 4% FA and processed as mentioned before.

### Bulk RNA sequencing of ablation experiment samples

Snap frozen cells were thawed into TRI reagent (sigma), RNA was isolated by Direct-zol RNA MiniPrep kit (Zymo research) according to the manufacturer instructions. RNA was processed by the mcSCRBseq protocol^46^ with minor modifications. RT reaction was applied on 10 ng of total RNA with a final volume of 10µl (1x Maxima H Buffer, 1mM dNTPs, 2µM TSO* E5V6NEXT, 7.5% PEG8000, 20U Maxima H enzyme, 1µl barcoded RT primer). Subsequent steps were applied as mentioned in the protocol. Library preparation was done using Nextera XT kit (Illumina) on 0.6ng amplified cDNA. Library final concentration of 2.2pM was loaded on NextSeq 500 (Illumina) sequencing machine aiming for 20M reads per sample. Raw files were converted to FASTQ files using bcl2fastq package, to obtain the UMI counts, fastq reads were aligned to the mouse reference genome (GRCm38.84) using zUMI packge^47^ with the following parameters RD1 16bp, RD2 66bp with a barcode (i7) length of 8bp. UMI counts were processed with the edgeR package^48^. Our analysis included four intestinal samples of DT-injected mice and four from mock-injected, 48 hrs after injection (Table S5). We first filtered the genes by expression to maintain only genes with expression greater than 10^−5^ of the summed UMI counts of the sample in at least 2 samples and next ran the calcNormFactors function with default settings. We used the “robust” option in the glmQLFit function to robustly estimate the QL dispersion and glmQLFTest with default parameters to compute differential gene expression with Benjamini-Hochberg correction for multiple hypotheses (Table S6). Fig. 6d shows the volcano plots of all genes with with maximal expression higher than 5*10^−6^ in at least one of the samples, as well as maximal zonation higher than 5*10^−6^ in Moor et al.^4^.

## Data availability

All data has been deposited in GEO with accession code GSE134479.

## Code availability

All codes used in this study will be available upon request.

## Acknowledgements

S.I. is supported by the Henry Chanoch Krenter Institute for Biomedical Imaging and Genomics, The Leir Charitable Foundations, Richard Jakubskind Laboratory of Systems Biology, Cymerman-Jakubskind Prize, The Lord Sieff of Brimpton Memorial Fund, the I-CORE program of the Planning and Budgeting Committee and the Israel Science Foundation (grants 1902/ 12 and 1796/12), the Israel Science Foundation grant No. 1486/16, the Broad Institute-Israel Science Foundation grant No. 2615/18, the Chan Zuckerberg Initiative grant No. CZF2019-002434, the European Research Council under the European Union’s Seventh Framework Programme (FP7/2007-2013)/ERC grant agreement number 335122, the Bert L. and N. Kuggie Vallee Foundation and the Howard Hughes Medical Institute (HHMI) international research scholar award. O.D.K. is supported by NIH R35-DE026602 and U01-DK103147.

## Author contribution

S.I., K.B.H., and M.S.C. conceived the study. K.B.H. designed and performed most of the experiments. H.M. and A.E.M. performed LCM experiments. K.B.H., N.W.C. and I.A. performed single cell RNAseq. K.B.H., M.R., L.F. and D.R.M. performed smFISH experiments. R.K.Z, D.C.A. K.B.H. performed the the *Lgr5*^*DTRGFP*^ mice experiments with supervision of O.D.K and F.J.D.S., S.I. and K.B.H. performed bioinformatics analysis. A.E. assisted with data analysis. S.I. and K.B.H. wrote the manuscript. S.I. and M.S.C. supervised the study. All authors discussed the results and commented on the manuscript.

## References

1. Flier, L. G. van der & Clevers, H. Stem Cells, Self-Renewal, and Differentiation in the Intestinal Epithelium. Annu. Rev. Physiol. 71, 241–260 (2009).

2. Barker, N. et al. Identification of stem cells in small intestine and colon by marker gene Lgr5. Nature 449, 1003–7 (2007).

3. Potten, C. S. Stem cells in gastrointestinal epithelium: numbers, characteristics and death. Philos. Trans. R. Soc. Lond. B. Biol. Sci. 353, 821–830 (1998).

4. Moor, A. E. et al. Spatial Reconstruction of Single Enterocytes Uncovers Broad Zonation along the Intestinal Villus Axis. Cell (2018) doi: 10.1016/j.cell.2018.08.063.

5. Beumer, J. et al. Enteroendocrine cells switch hormone expression along the crypt-to-villus BMP signalling gradient. Nat. Cell Biol. 20, 909–916 (2018).

6. Gehart, H. et al. Identification of Enteroendocrine Regulators by Real-Time Single-Cell Differentiation Mapping. Cell 176, 1158-1173.e16 (2019).

7. Roulis, M. & Flavell, R. A. Fibroblasts and myofibroblasts of the intestinal lamina propria in physiology and disease. Differentiation 92, 116–131 (2016).

8. Degirmenci, B., Valenta, T., Dimitrieva, S., Hausmann, G. & Basler, K. GLI1-expressing mesenchymal cells form the essential Wnt-secreting niche for colon stem cells. Nature 558, 449–453 (2018).

9. Shoshkes-Carmel, M. et al. Subepithelial telocytes are an important source of Wnts that supports intestinal crypts. Nature 557, 242–246 (2018).

10. Kinchen, J. et al. Structural Remodeling of the Human Colonic Mesenchyme in Inflammatory Bowel Disease. Cell 175, 372-386.e17 (2018).

11. Aoki, R. et al. Foxl1-expressing mesenchymal cells constitute the intestinal stem cell niche. Cell. Mol. Gastroenterol. Hepatol. 2, 175–188 (2016).

12. Ramilowski, J. A. et al. A draft network of ligand–receptor-mediated multicellular signalling in human. Nat. Commun. 6, 7866 (2015).

13. Vento-Tormo, R. et al. Single-cell reconstruction of the early maternal-fetal interface in humans. Nature 563, 347–353 (2018).

14. Tan, D. W.-M. & Barker, N. Chapter Three - Intestinal Stem Cells and Their Defining Niche. in Current Topics in Developmental Biology (ed. Rendl, M.) vol. 107 77–107 (Academic Press, 2014).

15. Itzkovitz, S. et al. Single molecule transcript counting of stem cell markers in the mouse intestine. Nat. Cell Biol. 14, 106–114 (2011).

16. He, X. C. et al. BMP signaling inhibits intestinal stem cell self-renewal through suppression of Wnt–β-catenin signaling. Nat. Genet. 36, 1117 (2004).

17. Naba, A. et al. The extracellular matrix: Tools and insights for the “omics” era. Matrix Biol. 49, 10–24 (2016).

18. Simon-Assmann, P., Spenle, C., Lefebvre, O. & Kedinger, M. The role of the basement membrane as a modulator of intestinal epithelial-mesenchymal interactions. Prog. Mol. Biol. Transl. Sci. 96, 175–206 (2010).

19. Meran, L., Baulies, A. & Li, V. S. W. Intestinal Stem Cell Niche: The Extracellular Matrix and Cellular Components. Stem Cells Int. 2017, (2017).

20. Karlsson, L., Lindahl, P., Heath, J. K. & Betsholtz, C. Abnormal gastrointestinal development in PDGF-A and PDGFR-α deficient mice implicates a novel mesenchymal structure with putative instructive properties in villus morphogenesis. Development 127, 3457–3466 (2000).

21. Cretoiu, D., Cretoiu, S. M., Simionescu, A. A. & Popescu, L. M. Telocytes, a distinct type of cell among the stromal cells present in the lamina propria of jejunum. Histol. Histopathol. 27, 1067–1078 (2012).

22. Greicius, G. et al. PDGFRα + pericryptal stromal cells are the critical source of Wnts and RSPO3 for murine intestinal stem cells in vivo. Proc. Natl. Acad. Sci. U. S. A. 115, E3173–E3181 (2018).

23. Stzepourginski, I. et al. CD34+ mesenchymal cells are a major component of the intestinal stem cells niche at homeostasis and after injury. Proc. Natl. Acad. Sci. U. S. A. 114, E506–E513 (2017).

24. Carmon, K. S., Gong, X., Lin, Q., Thomas, A. & Liu, Q. R-spondins function as ligands of the orphan receptors LGR4 and LGR5 to regulate Wnt/beta-catenin signaling. Proc Natl Acad Sci U A 108, 11452–7 (2011).

25. Tian, H. et al. A reserve stem cell population in small intestine renders Lgr5-positive cells dispensable. Nature (2011) doi: 10.1038/nature10408.

26. Biton, M. et al. T Helper Cell Cytokines Modulate Intestinal Stem Cell Renewal and Differentiation. Cell 175, 1307-1320.e22 (2018).

27. Zeisel, A. et al. Molecular Architecture of the Mouse Nervous System. Cell 174, 999-1014.e22 (2018).

28. Jaitin, D. A. et al. Massively Parallel Single-Cell RNA-Seq for Marker-Free Decomposition of Tissues into Cell Types. Science 343, 776–779 (2014).

29. Butler, A., Hoffman, P., Smibert, P., Papalexi, E. & Satija, R. Integrating single-cell transcriptomic data across different conditions, technologies, and species. Nat. Biotechnol. 36, 411–420 (2018).

30. Horiguchi, H. et al. ANGPTL2 expression in the intestinal stem cell niche controls epithelial regeneration and homeostasis. EMBO J. 36, 409–424 (2017).

31. Wei, X. et al. Spatially Restricted Stromal Wnt Signaling Restrains Prostate Epithelial Progenitor Growth through Direct and Indirect Mechanisms. Cell Stem Cell 24, 753-768.e6 (2019).

32. Kirsch, N. et al. Angiopoietin-like 4 Is a Wnt Signaling Antagonist that Promotes LRP6 Turnover. Dev. Cell 43, 71-82.e6 (2017).

33. Tetteh, P. W. et al. Replacement of Lost Lgr5-Positive Stem Cells through Plasticity of Their Enterocyte-Lineage Daughters. Cell Stem Cell (2016) doi: 10.1016/j.stem.2016.01.001.

34. Amin, N. & Vincan, E. The Wnt signaling pathways and cell adhesion. Front. Biosci. Landmark Ed. 17, 784–804 (2012).

35. Bengoa-Vergniory, N., Gorroño-Etxebarria, I., González-Salazar, I. & Kypta, R. M. A Switch From Canonical to Noncanonical Wnt Signaling Mediates Early Differentiation of Human Neural Stem Cells. STEM CELLS 32, 3196–3208 (2014).

36. Beyaz, S. et al. High-fat diet enhances stemness and tumorigenicity of intestinal progenitors. Nature 531, 53–58 (2016).

37. Yilmaz, Ö. H. et al. mTORC1 in the Paneth cell niche couples intestinal stem-cell function to calorie intake. Nature 486, 490–495 (2012).

38. Lee, J.-H. et al. Anatomically and Functionally Distinct Lung Mesenchymal Populations Marked by Lgr5 and Lgr6. Cell 170, 1149-1163.e12 (2017).

39. Moor, A. E. et al. Global mRNA polarization regulates translation efficiency in the intestinal epithelium. Science 357, 1299–1303 (2017).

40. Bray, N. L., Pimentel, H., Melsted, P. & Pachter, L. Near-optimal probabilistic RNA-seq quantification. Nat. Biotechnol. 34, 525–527 (2016).

41. Pimentel, H., Bray, N. L., Puente, S., Melsted, P. & Pachter, L. Differential analysis of RNA-seq incorporating quantification uncertainty. Nat. Methods 14, 687–690 (2017).

42. Preibisch, S., Saalfeld, S. & Tomancak, P. Globally optimal stitching of tiled 3D microscopic image acquisitions. Bioinforma. Oxf. Engl. 25, 1463–1465 (2009).

43. Schindelin, J. et al. Fiji: an open-source platform for biological-image analysis. Nat. Methods 9, 676–682 (2012).

44. Lyubimova, A. et al. Single-molecule mRNA detection and counting in mammalian tissue. Nat. Protoc. 8, 1743–1758 (2013).

45. Parekh, S., Ziegenhain, C., Vieth, B., Enard, W. & Hellmann, I. zUMIs - A fast and flexible pipeline to process RNA sequencing data with UMIs. GigaScience 7, (2018).

46. Bagnoli, J. W. et al. Sensitive and powerful single-cell RNA sequencing using mcSCRB-seq. Nat. Commun. 9, 2937 (2018).

47. Parekh, S., Ziegenhain, C., Vieth, B., Enard, W. & Hellmann, I. zUMIs - A fast and flexible pipeline to process RNA sequencing data with UMIs. bioRxiv 153940 (2018) doi: 10.1101/153940.

48. Robinson, M. D., McCarthy, D. J. & Smyth, G. K. edgeR: a Bioconductor package for differential expression analysis of digital gene expression data. Bioinformatics 26, 139–140 (2010).

